# Understanding the Sources of Performance in Deep Drug Response Models Reveals Insights and Improvements

**DOI:** 10.1101/2024.06.05.597337

**Authors:** Nikhil Branson, Pedro R. Cutillas, Conrad Bessant

## Abstract

Anti-cancer drug response prediction (DRP) using cancer cell lines plays a vital role in stratified medicine and drug discovery. Recently there has been a surge of new deep learning (DL) models for DRP that improve on the performance of their predecessors. However, different models use different input data types and neural network architectures making it hard to find the source of these improvements. Here we consider multiple published DRP models that report state-of-the-art performance in predicting continuous drug response values. These models take the chemical structures of drugs and omics profiles of cell lines as input. By experimenting with these models and comparing with our own simple benchmarks we show that no performance comes from drug features, instead, performance is due to the transcriptomics cell line profiles. Furthermore, we show that, depending on the testing type, much of the current reported performance is a property of the training target values. To address these limitations we create novel models (BinaryET and BinaryCB) that predict binary drug response values, guided by the hypothesis that this reduces the noise in the drug efficacy data. Thus, better aligning them with biochemistry that can be learnt from the input data. BinaryCB leverages a chemical foundation model, while BinaryET is trained from scratch using a transformer-type model. We show that these models learn useful chemical drug features, which is the first time this has been demonstrated for multiple DRP testing types to our knowledge. We further show binarising the drug response values is what causes the models to learn useful chemical drug features. We also show that BinaryET improves performance over BinaryCB, and over the published models that report state-of-the-art performance.

## 1 Introduction

Anti-cancer drug response prediction (DRP) has three main aims that, if delivered, would each improve patient outcomes or decrease treatment costs. These aims are to: (a) repurpose existing drugs, (b) tailor more effective treatments to specific groups or individuals and (c) help discover novel drugs. Cancer cell lines are at the heart of most DRP studies because they give a proxy for patient data while offering an abundance of publicly available drug screening data [1]. Several large-scale public databases provide drug response measurements and omics cell line profiles. These include the Genomics of Drug Sensitivity in Cancer (GDSC) [2], the cancer cell line encyclopedia (CCLE) and the Cancer Therapeutic Response Portal (CTRP) [3]. See [1, 4] for in-depth comparisons of the different databases. These databases typically contain hundreds of thousands of drug response measurements. For example, GDSC2 has 969 cell lines screened for up to 297 drugs (although not all cell lines have been screened for all drugs).

The availability of such datasets has resulted in a surge in new deep learning (DL) methods for predicting drug efficacy [5, 6, 7, 8, 9, 10, 11, 12]. The dominant paradigm for DRP models is to take omics profiles of cancer cell lines and drug structures as inputs to predict drug response. Such models have been reported to show state-of-the-art performance [7, 8, 9, 10]. See [12] for a review of DL methods in DRP. DRP models typically feed each of the two inputs (cell line and drug) through separate branches, to encode these inputs. They then use the fused encoded representations to predict how effective the input drug is for the input cell line. Figure 1 shows a diagrammatic representation of this. Earlier DRP models used the same subnetwork architectures for the drug and cell line branches, e.g. tCNNS used two subnetworks made from convolutional layers [13]. More recent models have used different subnetworks for the branches to leverage the different modalities of the input data.

**Figure 1:**
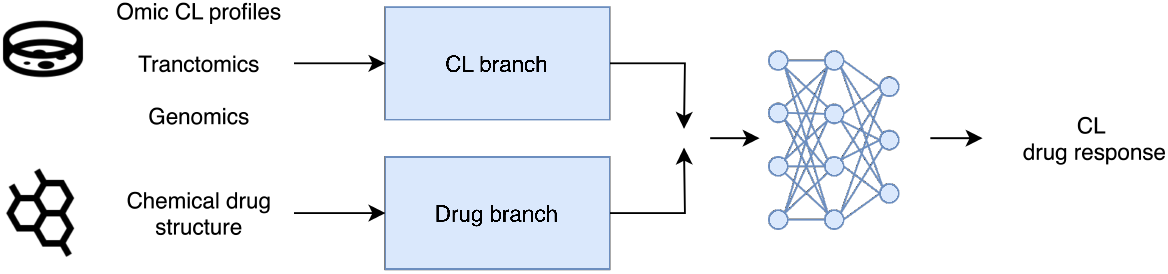
The general architecture of the models considered in this paper. Separate branches are used to encode the omics profiles and drug structures before being combined and fed through an MLP.

Typically DRP models have used convolutional or dense layers for the cell line branches [13, 7, 8, 9, 14, 15, 16, 17, 18]. In addition to these layers, there have been DRP models that use transformers [7], and graph convolutional layers [16, 9, 15] for their drug branches. There is a large variation in the architectures of the drug branches because drugs are three-dimensional chemical substances made of atoms bonded together. Therefore, there are many different ways to both numerically represent drugs and extract features from these representations. Due to the above, there are multiple differences in the representation of data and network structures of DRP models. Thus, when a new model outperforms an old one it is not clear if the improvement comes from enhancements to the model’s architecture, a better representation of the input data or a combination of the two. To compound this problem, DRP models are normally trained and tested in three different ways corresponding to the three different aims of DRP and it is standard for only one testing type to be used for ablation studies. Furthermore, the results and metrics can be stratified in two different ways, which we will show can also significantly impact the reported performance. However, this has been overlooked in previous studies.

Thus, in this paper, we look at how the different data types and subnetworks of three models, with reported state-of-the-art performance, impact their performance for DRP for the three testing types. These testing types are (1) mixed set (2) cancer blind and (3) drug blind testing where evaluation is done using unseen drug cell line pairs, cell lines and drugs respectively. The three models we use are tCNNS [13], DeepTTA [7], and GraphDRP [9]. Each of these models uses different drug branch architectures. tCNNS uses convolutional layers, DeepTTA uses a transformer and GraphDRP uses graph convolutional layers. We also introduce three simple but effective null hypothesis benchmarks, one for each testing type. These benchmarks do not use any omics data or drug structures. The benchmarks test how much performance improvement using omics data and drug structures adds and show how much performance can be attributed to patterns in the training truth values. This study shows that, for multiple testing types and models, all or most of the performance can be attributed to patterns in the training truth values. Furthermore, we find that, for multiple testing types, no performance comes from input chemical drug structures and instead, performance is due to the transcriptomics profiles.

For all of the above the models are trained and tested using continuous response values, as was done for the original implementations of these models [13, 7, 9] and is typical for DRP [17]. However, there is significant experimental noise in these response values due to the complex nature of the wet lab experiments [19, 20]. Thus, we hypothesise that the models are overfitting to the experimental noise in these values rather than learning chemically relevant features that are predictive of drug sensitivity and that, by binarising these response values, to remove some of the experimental noise the models can instead learn useful chemical features, resulting in better performance.

We test this by creating two novel transformer-based models BinaryET (encoder transformer), and BinaryCB (binary ChemBERTa) that predict binary drug response values. BinaryCB uses the chemical foundation model ChemBERTa [21, 22], for its drug branch. ChemBERTa is pre-trained using a large corpus of SMILES strings and has a BERT-like architecture [23]. In contrast, we train BinaryCB from scratch. We then conduct ablation studies on BinaryET and BinaryCB, removing the drug branch to show the models learn useful drug features. To our knowledge, this is the first time chemical drug structures have been shown to improve performance across multiple DRP testing types.

Comparing BinaryET with BinaryCB shows that the chemical foundational model does not improve performance. Re-training DeepTTA with binarised truth values we find that it can also learn useful chemical drug features, showing that binarising the drug response values is what causes the models to learn useful drug features. Furthermore, we show that BinaryET improves performance over the published models that report state-of-the-art performance. We highlight our main contributions below:

- For mixed set testing we find that our null hypothesis benchmark outperforms the published models with reported state-of-the-art performance thus, performance can be explained entirely by patterns in the truth values.
- We show that for cancer blind testing, no performance is due to the drug structures instead performance is driven by transcriptomics profiles, with metric stratification significantly impacting results.
- We introduce BinaryET and BinaryCB models predicting binary drug response values, hypothesising reduced experimental noise and improved alignment with biochemistry. BinaryCB employs ChemBERTa, a RoBERTa-like chemical foundation model, while BinaryET uses transformer encoder layers trained from scratch.
- We find that BinaryET and BinaryCB effectively leverage chemical drug structures and that binarizing response values are key to learning useful drug features.
- We find BinaryET outperforms BinaryCB and the SOTA models re-trained with binary response values.

## 2 Related Work

Chen and Zhang [24] recreated a number of DL DRP models to compare performance. Li et al. [25] investigate interpretable DL DRP models to see if interpretability comes at the cost of performance. Li et al. also show the strong performance of null hypothesis benchmarks compared to the DL models. However, due to the challenging nature of recreating models from disparate codebases and implementations there is limited work that involves previously published DRP models [12].

Prior work that aims to understand deep learning models in other applications has provided many invaluable insights and improvements. These insights come from an understanding of how the models work[26, 27, 28, 29, 30]. But also from an understanding of the metrics used to evaluate the models [31, 32].Where it is vital that the metrics track what researchers are trying to measure and reflect the applications of the models in practice. Thus, there is significant potential and scope to explore DL DRP models and the source of their performance, both in terms of how the metrics are calculated and the models themselves.

Large models pre-trained using chemical drug structures have shown strong performance and have been observed to exhibit scaling laws [33, 21, 22, 34] leading some researchers to name them chemical foundation models [21, 35]. These models have been utilised for many downstream tasks [36, 21, 22, 34]. However, to our knowledge, they have not yet been applied in DRP.

## 3 Methods

To make the definition of DRP more concrete consider 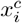 and 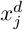 representation of the *i*^*th*^ cell line and *j*^*th*^ drug respectively and associated truth value *y*_*i,j*_. Here *y*_*i,j*_ is the drug response value associated with the (*i, j*) cell line drug pair, which describes how effective the *j*^*th*^ drug is at killing the *i*^*th*^ cell line. For example, *y* could be an IC50 value, the concentration of a drug needed to reduce the activity of a cell line by 50%.

A DRP model, *M* created by a learning algorithm takes 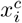 and 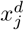 as inputs and predicts for the corresponding truth value 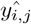 such that 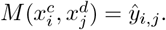.

### 3.1 Evaluating DRP models

The aim of drug response prediction is to find a model using *T* that performs well on unseen data. This can be done by evaluating the model on some held-out testing data *E*. There are three common ways of constructing *T* and *E* corresponding to the three different objectives for DRP as outlined in the introduction.

1. Mixed-set: any drug and any cell line can be in either *T* or *E*, but a drug cell line pair can only be in one set.
2. Cancer-blind: cell lines in *T* can not be in *E*, but all drugs are be in both sets.
3. Drug-blind: drugs in *T* can not be in *E*, but all cell lines are in both either set.

Figure 2 diagrammatically shows these different splits.

**Figure 2:**
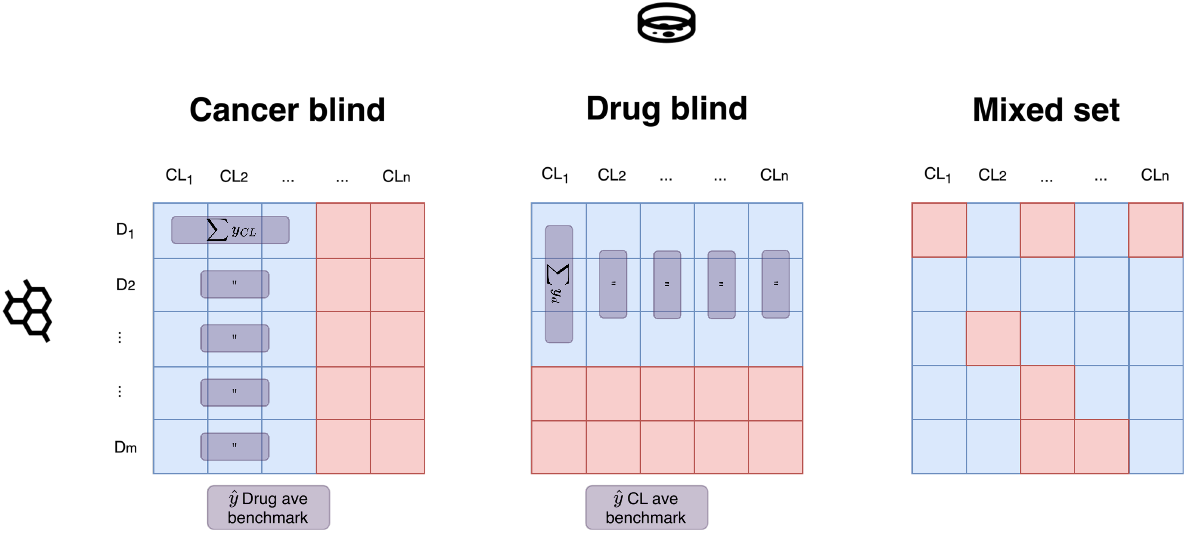
The different ways of splitting response values. Each square represents the efficacy of drug *j* for cell line *i*. The blue and red squares represent training and testing response values respectively. The input drug and cell line representations are also split into the appropriate set, given by the response. The purple boxes represent how the predictions for the null benchmarks introduced in section 3.4 are calculated.

Using mixed-set testing shows if the model could be used for drug repurposing. This is because, for a given dataset a cell line in this dataset is typically not evaluated against all drugs in the dataset. Therefore, the model can be used to evaluate drug cell line pairs that have not yet been explored. This is a quick and inexpensive way to predict if a known drug (drug in the training set) can be repurposed for a known cancer subtype. Cancer-blind testing shows if the model could be used to find candidate drugs that might be effective for cancer sub-types, not in the training set. This would be useful in drug discovery, where cancer cell lines can be used to narrow down candidate drugs. Furthermore, cancer-blind testing is a suitable method for simulating how good a model would be in a stratified medicine context. Where the model would need to predict efficacy for samples it was not trained on. For example to predict if a drug, or set of drugs, is suitable for new patients. A model that performs well using drug-blind testing could be used to find novel anti-cancer drugs from just its structure. This is because the model would be able to predict the responses of a candidate drug to known cell lines.

### 3.2 Stratification of results in metric calculations

There are two inputs to DRP models. Thus, metrics can be stratified in two different ways, by cell line or by drug. We consider cancer blind testing in the following to make our examples concrete. We define stratifying results by cell line as finding a metric for each cell line in the test set individually, across all drugs, before averaging the results. Similarly, we define stratifying the results by drug as finding a metric for each drug, across all cell lines in the test set, before averaging the results. In contrast, a non-stratified metric is found once across all cell line drug pairs in the testing set without averaging. Thus, there are three different ways performance metrics can be reported. In DRP studies this is typically not discussed and only stratification by cell line is reported. However, as we will show, how results are stratified makes a big impact on cancer blind testing results.

The two different methods for stratification correspond to evaluating the model for distinct problems the model could be applied to. Stratifying by cell line corresponds to simulating the performance in a clinical stratified medicine context which aims to recommend a drug out of all the drugs the model is trained on, for a new patient. A model that performs well using this testing would also be directly useful in a drug discovery context where cell lines are used to narrow down candidate drugs. Specifically, the model could be used, to rank treatments from the set of all drugs the model has been trained on for an unseen cancer sub-type.

In contrast, stratifying by drug corresponds to simulating the performance in a clinical stratified medicine context where there is a set of new patients and you want to recommend if a given drug should be taken for each of these patients. This is typically the scenario in which DRP models are applied to [11, 10, 37, 38]. The corresponding application in a drug discovery context directly using cell lines, is where you have a candidate drug that has already been screened for some cell lines and you can then use the model to screen a set of many unseen cancer sub-types for efficacy. We provide the equations for calculating the stratified metrics in Appendix B.

### 3.3 Datasets and models

At a high level, our model, BinaryET consists of three subnetworks: a drug and cell line branch and a regressor, as shown in figure 1. Each branch separately encodes the input data before the outputs of the branches are concatenated and passed through a regressor. For our cell line branch, we used all the transcriptomic features for our cell line profiles as input to a three-layer multilayer perceptron (MLP). For our drug branch, we used SMILES strings tokenized with the ChemBERTa tokenizer [22] implemented in Hugging Face [39]. We then used the tokenized SMILES strings as input to a stack of transformer encoder layers [40]. We then passed the concatenated outputs of the two branches through a 3-layer MLP, that predicted the binarised response value of the input drug cell line pair. For BinaryCB we replaced the drug branch of BinaryET with ChemBERTa and fine-tuned the whole model during training. For further details of the datasets used and how we binarised the response IC50 values see Appendix D. For model hyperparameters see Appendix G.

An 80%, 10%, 10% train, validation, and test split was used for all testing and the validation data was used to select hyperparameters and the optimal number of epochs. We ran the models for three train test splits and for each split, we ran the model for three different seeds. We also repeated this for each of the testing types, mixed set, cancer blind and drug blind. This same training protocol was also used for the published models that we recreated. See Appendix E for further details on how we recreated these models to predict both continuous and binary response values.

### 3.4 Null Hypothesis Benchmarks

We create three null hypothesis benchmarks, (1) drug average (2) cell line (CL) average and (3) marker benchmark, for use in cancer blind, drug blind and mixed set testing respectively. None of these benchmarks use any omics data or drug structures. We note that the drug and CL average benchmarks have recently been shown to have strong performance [14, 25] but have not yet been fully explored. The benchmarks are defined as follows:

#### Drug average benchmark

predictions for response values for a given drug in the test set were simply calculated as the mean of all response values for that drug in the training set. This is shown in figure 2 under the cancer blind split. The figure shows that for a given drug, a row in the figure, the average is calculated across cell lines in the training set (the boxes in blue). This average is then the prediction for the cell line drug pairs in the test set for that drug (the red boxes). Thus, for the drug average benchmark the prediction for the drug, *d*, and any cell line, 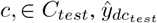 is given by

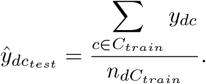

The sum runs over all cell lines in the training set *c ∈ C*_*train*_ where drug cell line pair (*d, c*) has an associated response value 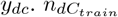 is the number of drug cell line pairs in the training set that include *d*. Note that due to missing response values 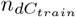 can be different for different drugs.

#### CL average benchmark

the above was done with cell lines instead of drugs. Thus, the prediction for the cell line, *c* and any drug in the testing set *d ∈ D*_*test*_ is given by

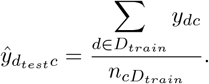

Where 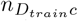 is the number of drug cell line pairs in the training set that include *c*. Similarly Figure 2 shows the predictions of the CL average benchmark under the drug blind split.

#### Marker benchmark

we define a marker representation of either a cell line or drug to mean a vector that uniquely identifies that cell line or drug but doesn’t have any biological or chemical properties. Here, we use a column vector to achieve this, by one-hot encoding each drug or cell line. The marker benchmark took cell line and drug inputs as marker representations and fed them through an MLP.

We hypothesised each benchmarks were effective for similar reasons:

- Drug average benchmark: there are drugs that are generally effective at killing most cell lines and other drugs that are ineffective for most cell lines.
- CL average benchmark: there are cell lines that are generally harder or easier to kill for most drugs.
- Marker benchmark: all drugs and cell lines are in the training set for mixed-set evaluation. Thus, a combination of the above properties can be learnt during training for inference.

Importantly, all of the null hypothesis benchmarks allowed us to see if adding omics data or drug structures improves performance and how much of model performance can be attributed to the above property of the data.

## 4 Results and Discussion

### 4.1 Cancer blind testing with continuous drug response values

Table 1 shows the results of cancer blind testing for the published model’s drug average benchmark and DeepTTA-DB. We created DeepTTA minus drug branch (DeepTTA-DB) to see how much impact chemical drug structures have on cancer blind performance. For DeepTTA-DB instead of DeepTTA’s transformer drug branch, we simply feed in a one-hot encoded marker representation of the drugs. See appendix C for details of the metrics we report. The reported performance difference seen between the stratified and non-stratified results (defined in section 3.2) shows that cell line stratification is an easier problem. This is expected and is due to the statistics of the response values specifically due to observation 1:

**Table 1:** Metrics for cell line and drug stratified cancer blind testing, for three models from the literature: tCNNS, DeepTTA, GraphDRP, DeepTTA-DB (a model that we create), and our null hypothesis drug average benchmark. See appendix C for details of the metrics we report. The best model performance per metric is highlighted in **bold** for this and the following tables. The uncertainties in this and all tables are the standard deviations across three model seeds.

| Method | MSE<br>CL strat | Pear<br>CL strat | R2<br>CL strat | Pear<br>drug strat | R2<br>drug strat |
| --- | --- | --- | --- | --- | --- |
| tCNNS | 2.41 ± 0.04 | 0.870 ± 0.003 | 0.61 ± 0.01 | 0.16 ± 0.02 | -0.21 ± 0.01 |
| GraphDRP | 2.41 ± 0.08 | 0.872 ± 0.004 | 0.60 ± 0.01 | 0.220 ± 0.009 | -0.26 ± 0.06 |
| Drug Average | 2.190 | 0.884 | 0.640 | N/A | -0.012 |
| DeepTTA-DB | 1.575 ± 0.009 | 0.902 ± 0.001 | **0.743 ± 0.004** | **0.526 ± 0.009** | **0.257 ± 0.004** |
| DeepTTA | **1.57 ± 0.02** | **0.904 ± 0.002** | **0.743 ± 0.005** | 0.523 ± 0.006 | 0.253 ± 0.008 |

#### Observation 1.

*Much of the variance in the response values between drugs can be explained by the average behaviour of the drugs*.

We note that Observation 1 is a direct implication of the hypothesis made in section 3.4, that there are drugs that are generally effective or ineffective. The drug average benchmark clearly demonstrates Observation 1 as it is directly built to reflect this property. Thus, when we test across all of the drugs, Observation 1 boosts the performance of the models. In contrast with drug stratification where performance is evaluated for a given drug, the variance cannot be explained by Observation 1 as we are looking across cell lines for that drug. This is again shown by the drug average benchmark that predicts the same value, for a given drug across all cell lines, and thus, gives an *R*^2^ *∼* 0 and has an undefined Pearson correlation coefficient. This is also why the models have a much better performance differential compared with the benchmark, for drug than cell line stratification. This shows the importance of considering how results are stratified, whereas DRP studies typically only report cell line stratified metrics and to our knowledge do not consider drug stratification. Furthermore, it shows that deep learning models can have the most impact for a drug stratified use case. Interestingly previous work using learning curves found that the performance of the drug average benchmark plateaued at larger dataset sizes but models using omics data did not. This suggests that the performance differential to the benchmark will increase as more data is collected [14].

Table 1 also shows the drug average benchmark outperforms both of the models that use genomic cell line profiles, tCNNS and GraphDRP, in terms of all metrics bar drug stratified Pearson correlation. Thus, these models are not generally learning useful features above those that can be derived from Observation 1. On the other hand, the two models that use transcriptomic cell line profiles, DeepTTA and DeepTTA-DB comfortably outperform the benchmark for all metrics. This suggests that transcriptomic profiles play a key role in cancer blind testing. To further explore this result we constructed tCNNS_Tran and GraphDRP_Tran, where we replaced the genomic cell line profiles of tCNNS and GraphDRP with transcriptomic profiles with all genes as features. The results for these models are shown in Table 12 in Appendix H. The table shows that tCNNS_Tran and GraphDRP_Tran both outperform the benchmark further showing that the strong performance is caused by the transcriptomic cell line profiles. There is also a strong theoretical justification for this result. Many of the processes in cancer can not be explained by genomic profiles (mutation and copy number variation data) alone [41]. In contrast, transcriptomics operates at the level of post-transcription so contains vital information about biochemical pathways, important for understanding diseases, that genomics does not [42]. Therefore, transcriptomics describes parts of the biology of cancer that genomics can not.

Table 1 also shows that DeepTTA-DB and DeepTTA perform equally within the margin of uncertainty. Therefore, the performance is due to the omics branch, further confirming that the transcriptomic cell line profiles cause the performance improvement seen over the drug average benchmark. We further explore this result by removing the drug branch from tCNNS_Tran and GraphDRP_Tran to create tCNNS_Tran-DB and GraphDRP_Tran-DB. As with DeepTTA-DB, tCNNS_Tran-DB and GraphDRP_Tran-DB are tCNNS_Tran and GraphDRP_Tran with a one-hot encoded marker representation of the drugs used in place of the respective drug branches. The results for these models are shown in Table 12 in Appendix H. The table shows that removing the drug branch from both models does not impact performance. This supports the above discussion, that the performance improvements over the benchmark are due to the transcriptomic cell line profiles and the models do not learn useful representations from the chemical drug structures.

We note that it is expected that for cancer blind testing transcriptomics profiles will contribute more than chemical drug profiles. This is because, in cancer blind testing, all drugs are in both the training and testing set. Therefore, the model can learn the distribution of the drug’s efficacy during training and directly use this information in the test set. This is demonstrated by the drug average benchmark that shows good performance, on the test set, only using the distribution of the drug efficacy in the train set. In contrast, the cell lines tested on are unseen, therefore the omics profiles are required for the model to discriminate on. This is because, for a given drug, there is no other way for the model to differentiate its prediction than with the cell line profiles.

We repeated the above analysis for two further test train splits. The values of all metrics are similar to those in table 1. Furthermore, the results agree with the conclusions arrived at in the above discussion. The results for two other test train splits are shown in Appendix H. We also experiment with first scaling the truth values to be between zero and one using the same method as in [13] before re-training and testing the models and found consistent results (Table 13 in Appendix H) Thus, showing the robustness of our results to different methods of scaling the response values.

### 4.2 Mixed set and drug blind testing with continuous drug response values

Table 2 shows the results of mixed set testing for the three published models and the marker benchmark. It shows that none of the published models outperform the marker benchmark, for all metrics. The marker benchmark does not use omics data or chemical drug features. This means the performance of the model comes from the truth values in the training set. Therefore, the published models are unable to use omics or drug data to improve performance over what information can be gained from the truth values in the training set. The marker benchmark is so successful because, in mixed set testing, every drug and cell line in the test set is also in the training set while unique drug cell line pairs are only in the training or testing set. Therefore, the model has ample information on the distribution of response values for any cell line or drug in the training set to learn from. Thus, rather than having to learn important cell lines or drug features, it can just use these distributions for predictions. We repeated the above analysis for two further test train splits to increase the reliability of our results. The results for these splits support Table 2 and are shown in Appendix I. We also note that DRP studies typically do their ablation studies for this testing type, limiting their usefulness in light of the above.

**Table 2:** Metrics for mixed set testing for three models from the literature and our null hypothesis marker benchmark. The published models do not outperform the benchmark for any metric. Note Transcript and mol graph are used as shorthand for transcriptomic and molecular graph respectively.

| Method | Drug branch | Omics branch | MSE | Pear | R2 |
| --- | --- | --- | --- | --- | --- |
| tCNNS | SMILES | Genomics | $1.256 \pm 0.004$ | $0.9104 \pm 0.0003$ | $0.8283 \pm 0.0005$ |
| GraphDRP | Mol graph | Genomics | $1.06 \pm 0.04$ | $0.930 \pm 0.001$ | $0.856 \pm 0.005$ |
| DeepTTA | SMILES | Transcript | $0.98 \pm 0.01$ | $0.931 \pm 0.001$ | $0.866 \pm 0.002$ |
| Marker Benchmark | Marker | Marker | **$0.88 \pm 0.01$** | **$0.9379 \pm 0.0007$** | **$0.880 \pm 0.001$** |

To further investigate this result we created marker versions of each of the literature models. We did this by replacing the drug branches and omics inputs of the models with a marker representation before re-training and testing the models. The results given in Appendix I show that all marker versions of the models outperform the original models. This supports the hypothesis that the models are learning the distribution of efficacy/susceptibility of drugs/cell lines by associating the IC50 values with a marker representation of the inputs, instead of learning biological or chemically relevant features from the input omics or chemical structures and removing the omics/chemical input features allows the model to more easily do this. These results show the importance of using marker representations to benchmark DRP models for mixed set testing.

For drug blind testing we found a much greater variation in performance between stratified and non-stratified testing than we found for cancer blind testing. There was also a greater variation between different train test splits than we found for the other testing types. The results for all splits, stratified and non-stratified testing are shown in Appendix J. These results show that while there are large differences between the test train splits, for each split there are metrics that outperform the CL average benchmark. Therefore, the omics and drug structures can provide a benefit for non-stratified drug blind testing. However, more needs to be done to improve the model’s drug blind testing abilities so that they outperform the benchmark consistently by improving model stability.

### 4.3 Evaluation of BinaryET

Table 3 shows the results for drug stratified cancer blind testing using binary response values. It shows our model, BinaryET outperforms all other models that had previously reported state-of-the-art performance, across all metrics. All models outperform the drug average benchmark for AUC and AUPR. We note that by definition the drug average benchmark has an AUC of 0.5, as is seen in table 3. The results for two further train test splits are shown in Appendix K, they support the results in table 3. Appendix K also shows the metrics for the same set of results but with cell line stratification and without stratification. It supports the result from table 3 that BinaryET outperforms all other models. However, unlike in table 3 the drug average benchmark outperforms GraphDRP and tCNNS for both AUC and AUPR. These results also support the difference between cell line and drug stratified metrics seen for continuous response values.

**Table 3:** Metrics for drug stratified drug blind testing for BinaryET (ours) compared with the published models, retrained to predict binary response values, and the null hypothesis drug average benchmark.

| Method | AUC | AUPR |
| --- | --- | --- |
| Drug Average | 0.500 | 0.320 |
| tCNNS | $0.598 \pm 0.008$ | $0.40 \pm 0.01$ |
| GraphDRP | $0.6173 \pm 0.0009$ | $0.421 \pm 0.007$ |
| DeepTTA | $0.747 \pm 0.008$ | $0.53 \pm 0.02$ |
| BinaryET | **$0.771 \pm 0.003$** | **$0.569 \pm 0.004$** |

Table 4 shows the results for the ablation study where we remove the drug branch from BinaryET. The table shows for all splits and ways of stratifying the results removing the drug branches decreases the performance of our model. Therefore, BinaryET is able to extract useful representations from the chemical drug structures. Appendix K shows this result also holds for DeepTTA. This is in contrast to the results we found when continuous truth values are used (Table 1). Thus, binarising the truth values allows the models to successfully leverage chemical drug structures. We hypothesise that this is because due to the experimental noise in the continuous response values, they do not fully reflect the underlying causal biochemistry of the experiment. Thus, there are not chemically relevant features that are predictive of continuous drug sensitivity that the models can learn, over what can be learnt from the transcriptomics. Then, binarising these response values removes some of the experimental noise better aligning the binary response values with underlying biochemistry, which the input chemical drug structures can explain allowing models to learn useful features from these inputs.

**Table 4:** Ablation study for BinaryET removing the drug branch BinaryET-DB, for all types of stratified cancer blind testing.

| Stratification type | BinaryET<br>AUC | BinaryET-DB<br>AUC | BinaryET<br>AUPR | BinaryET-DB<br>AUPR |
| --- | --- | --- | --- | --- |
| Drug Strat | **$0.771 \pm 0.003$** | $0.76 \pm 0.01$ | **$0.569 \pm 0.004$** | $0.55 \pm 0.02$ |
| Cell Line Strat | **$0.9305 \pm 0.0006$** | $0.928 \pm 0.001$ | **$0.822 \pm 0.002$** | $0.817 \pm 0.001$ |
| No Strat | **$0.9158 \pm 0.0006$** | $0.913 \pm 0.001$ | **$0.8084 \pm 0.0002$** | $0.8033 \pm 0.0008$ |

Table 5 shows the results for BinaryCB, where we have replaced the drug branch in BinaryET with ChemBERTa. We have evaluated different versions of ChemBERTa pre-trained using databases of SMILES strings of different sizes, from 40, 000 SMILES strings from the ZINC database to 77, 000, 000 from the PubChem database. The table also shows the results for an ablation of BinaryCB, Marker DB where we replace the drug branch with a marker drug input. The table also includes results for a model with the same architecture and hyperparameters as BinaryCB but without any pre-training (No PT) which is the same as BinaryET with different hyperparameters. The tables show that while BinaryCB does learn useful chemical drug features, the pre-training does not provide any performance improvements over training a model from scratch. The table also shows that increasing the pre-training size of ChemBERTa does not improve performance. Furthermore, moving from ZINC to PubChem leads to a drop in performance despite an order of magnitude increase in the number of SMILES strings used for pre-training. A possible reason for this is that the distribution of SMILES strings taken to train ChemBERTa from PubChem may be further away from anticancer drugs than those that were chosen from ZINC.

**Table 5:** Pefomrance of BinaryCB with different versions of ChemBERTa used for the drug branch. Performance is the average of three model seeds on the validation set.

|  | BCE (Loss) | AUC | AUPR |
| --- | --- | --- | --- |
| zinc40k | $0.3448 \pm 0.0003$ | $0.9169 \pm 0.0002$ | $0.8561 \pm 0.0002$ |
| zinc100k | $0.345 \pm 0.001$ | $0.9170 \pm 0.0006$ | $0.8558 \pm 0.0006$ |
| zinc250k | $0.3432 \pm 0.0006$ | $0.9169 \pm 0.0002$ | $0.8564 \pm 0.0003$ |
| Pub5M | $0.354 \pm 0.001$ | $0.9140 \pm 0.0005$ | $0.8512 \pm 0.0005$ |
| Pub10M | $0.357 \pm 0.003$ | $0.914 \pm 0.002$ | $0.851 \pm 0.002$ |
| Pub77M | $0.360 \pm 0.001$ | $0.9123 \pm 0.0006$ | $0.849 \pm 0.001$ |
| Marker DB | $0.3502 \pm 0.0007$ | $0.9136 \pm 0.0001$ | $0.8506 \pm 0.0005$ |
| No PT | $0.3429 \pm 0.0007$ | $0.917 \pm 0.001$ | $0.857 \pm 0.001$ |
| BinaryET | **$0.342 \pm 0.002$** | **$0.919 \pm 0.001$** | **$0.858 \pm 0.002$** |

Table 6 shows the results for mixed set testing with binary response values. It shows that our model, BinaryET, outperforms all other models that had previously reported state-of-the-art performance. In contrast to the continuous case, all models outperform the null hypothesis benchmark. However, it still performs comparatively to the models suggesting that much of the performance is due to patterns directly inferred from the training truth values. Furthermore, the table shows that removing the drug branch from BinaryET decreases the performance. Thus, BinaryET also learns useful information from its drug branch for mixed-set in addition to cancer blind testing. The results for two further train test splits are shown in appendix L, they agree with the results in table 6.

**Table 6:** Metrics for mixed-set testing for BinaryET (ours) compared with the published models, retrained to predict binary response values, and the null hypothesis marker average benchmark.

|  | AUC | AUPR |
| --- | --- | --- |
| Marker benchmark | $0.9276 \pm 0.0001$ | $0.8634 \pm 0.0001$ |
| tCNNS | $0.928 \pm 0.001$ | $0.863 \pm 0.001$ |
| GraphDRP | $0.9376 \pm 0.0007$ | $0.882 \pm 0.001$ |
| DeepTTA | $0.9435 \pm 0.0008$ | $0.892 \pm 0.001$ |
| BinaryET-DB | $0.9458 \pm 0.0006$ | $0.896 \pm 0.001$ |
| BinaryET | **$0.9470 \pm 0.0004$** | **$0.8983 \pm 0.0008$** |

## 5 Conclusion

In this paper we recreated three drug response prediction (DRP) models, that have reported state-of-the-art performance. We also created null hypothesis benchmarks that did not use omics data or chemical features to help understand the published model’s sources of performance. These benchmarks showed strong performance relative to the models suggesting that they should be used when evaluating future DRP research. They also revealed that for multiple testing types, the performance could partially or fully be explained by patterns in the training truth values. For cancer blind testing we found, using ablation studies, that none of the model performance comes from their chemical drug structures, instead, it is due to the transcriptomics cell line profiles. To address these limitations we created BinaryET and BinaryCB, to predict binary drug response values guided by the hypothesis that this will remove some of the experimental noise allowing them to learn useful chemical features. We find this to be the case for multiple testing types. Furthermore, we show BinaryET improves upon the performance of the models that have reported state-of-the-art performance.

## Acknowledgments and Disclosure of Funding

This work was supported by PhD studentship funding from the Wellcome Trust [218584/Z/19/Z]

## A Appendix / supplemental material

### B Calculating stratified metrics

For a cell line stratified metric *MC*_*strat*_,

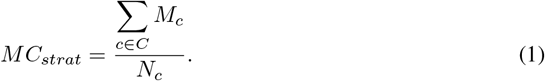

Where the sum runs over, *C* all cell lines in the test set and *N*_*c*_ is the number of cell lines in the test set. *M*_*c*_ is the metric *M* for cell line *c* such that

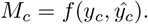

Where *y*_*c*_ and *ŷ*_*c*_ are the truth values and predicted values for the cell line drug pairs that include *c* respectively. *f* is a function that gives metric *M* for example it could be the mean squared error. In contrast drug-stratified metric is instead given by *MD*_*strat*_,

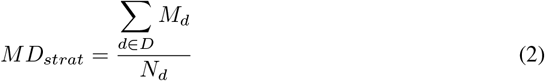

Where the sum runs over, *D* all drugs and *N*_*d*_ is the number of drugs. *M*_*d*_ is the metric *M* for drug *d* such that

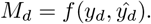

Where *y*_*d*_ and *ŷ*_*d*_ are the truth values and predicted values for all drug cell lines pairs that include *d*, and are in the test set.

Similarly, drug stratified drug blind testing is defined by equation 2 but with the sum only running over drugs in the testing set, and *N*_*d*_ giving the number of drugs in the testing set. Furthermore, cell line stratified drug blind testing is defined by equation 1 but with the sum running over all cell lines, and where *y*_*c*_, *ŷ*_*c*_ are only the truth and predicted values for the drug cell line pairs in the test set that contain *c*.

A non-stratified *M* is simply given by

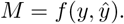

Where *y* and *ŷ* are the truth values and predicted values for the cell line drug pairs in the test set respectively.

### C Metrics reported

When evaluating models that predicted continuous drug response values we calculate Mean squared error (MSE), Pearson correlation coefficient (Pear) and the Coefficient of determination (R2). We report Pear and R2 for both cell line (CL) stratification and drug stratification as well as non-stratification, as described above.

When evaluating models that predicted binary drug response values we calculate the area under the receiver operating characteristic curve (AUC) and the area under the precision-recall curve (AUPR).

### D Dataset details

Across this study, we used transcriptomic cell line profiles from the genomics of drug sensitivity in cancer database (GDSC) [2]. For drug response data we used IC50 values from GDSC2, where IC50 is the standard measure of drug response and was used in the original papers for tCNNS DeepTTA and GraphDRP. These downloaded IC50 values were continuous measurements. Thus, we binarised them for the second part of our study when considering binary drug efficacy. We used the same method to binarise the IC50 values as Liu et al. [8], where a drug is considered ineffective if its IC50 value is more than the maximum concentration used during screening. We downloaded SMILES representations of the drugs from PubChem [43].

Table 7 shows the full dataset size used when training and testing the modles for both binary and continuous IC50 values. Only cell lines that had IC50 values and both genomics and transcriptomics cell line profiles were kept. Four drugs were removed when binarising IC50 values as they had multiple maximum concentration values. We note that not all drug cell line pairs have IC50 values in GDSC2. Thus, we also removed drug cell line pairs without IC50 values, hence why we have less than 163, 624 and 160, 008 drug cell line pairs for continuous and binary IC50 values respectively.

**Table 7:** Dataset size for binary and continuous IC50 values. An %80, %10, %10 train validation test split was used. For drug-stratified binary cancer blind splitting the number of drugs we tested was 147, 153, 148 for train test split 1, 2, and 3 respectively, due to AUC and AUPR not being defined for the drugs removed.

|  | Number of<br>cell lines | Number of<br>drugs | Number of<br>drug cell line pairs |
| --- | --- | --- | --- |
| Continuous | 904 | 181 | 147,713 |
| Binary | 904 | 177 | 144,101 |

**Table 8:** Number of positive and negative examples after data splitting for binary response values.

|  | S1 Negative | S1 Positive | S2 Negative | S2 Positive | S3 Negative | S3 Positive |
| --- | --- | --- | --- | --- | --- | --- |
| train | 79359 | 35910 | 79448 | 35898 | 79439 | 36013 |
| test | 10637 | 4047 | 10015 | 4394 | 10151 | 4230 |
| val | 9339 | 4809 | 9872 | 4474 | 9745 | 4523 |

### E Details of published models used

For genomics cell line profiles we used genomics data from the genomics of drug sensitivity in cancer database (GDSC) [2]. Genomics profiles that included, genetic mutation and copy number variations information, were used to recreate tCNNS and GraphDRP. For transcriptomic profiles, we used transcriptomics data from GDSC. Transcriptomics profiles were used to recreate DeepTTA. We only kept cell lines that were in all three of the above datasets. Thus, 904 cell lines were used in the following analysis.

The omics data in GDSC has already undergone standard preprocessing. Where the transcriptomics data is preprocessed using the robust multi-array analysis algorithm (RMA) [44]. This is the same data that was used in DeepTTA. Similarly, we directly used the binary genomics data from GDSC as was done in tCNNS and GraphDRP. Furthermore, the IC50 values provided by GDSC are natural logarithm transformed.

SMILES (simplified molecular-input line-entry system) and molecular graphs were used as the drug representations with chemical properties. SMILES were used in the tCNNS and DeepTTA models, while molecular graphs were used in the GraphDRP model. In a SMILES representation, each molecule is represented as a string of characters. In molecular graph representations of molecules/drugs, each node represents an atom in the molecule and each edge represents a bond between the atoms. Only drugs with SMILES strings were kept, leading to us using 181 drugs for the following analysis.

When retraining and testing these models with binary response values we used binary cross entropy for the loss function.

### F Hardwear

Models were trained using nvidia A100 GPUs, each model was trained on one GPU.

### G Model hyperparameters

The model hyperparameters for BinaryET are shown in table 9.

**Table 9:** BinaryET hyperparameters the first 6 parameters refer to the drug branch (Transformer encoder layers).

| Hyperparameters | Value |
| --- | --- |
| Nb transformer (TF) encoder layers | 8 |
| Nb attention heads | 8 |
| TF feed forward dim | 2048 |
| Embedding dimension | 128 |
| Dropout | 0.01 |
| layer_norm_eps | 1e-05 |
| Activation | relu |
| Nb classifier nodes layer 1 | 1024 |
| Nb classifier nodes layer 2 | 256 |
| Nb classifier nodes layer 3 | 64 |
| Loss | binary cross entropy |
| Batch size | 128 |
| Peak learning rate | 5e-05 |
| Optimiser | AdamW |
| learning rate scheduler | cosine with warmup |
| epochs | 100 |
| Nb warm up learning rate steps | 384 |

For the marker benchmark, 3 dense hidden layers were for the MLP with 4096 neurons per layer, with ReLU activation, followed by an output dense layer with one node.

### H Cancer blind testing

This section shows the tables for additional cancer blind testing with continuous response values. Table 10 shows two additional train test splits for tCNNS, GraphDRP, DeepTTA-DB and DeepTTA.

**Table 10:** Metrics for cancer blind testing for the second and thrid test train split.

| Split | Method | MSE | Pear<br>CL strat | R2<br>CL strat | Pear<br>drug strat | R2<br>drug strat | MSE<br>drug strat |
| --- | --- | --- | --- | --- | --- | --- | --- |
| 2 | tCNNS | 2.4 ± 0.005 | 0.864 ± 0.002 | 0.620 ± 0.004 | 0.21 ± 0.04 | -0.12 ± 0.03 | 2.45 ± 0.03 |
|  | GraphDRP | 2.45 ± 0.05 | 0.867 ± 0.002 | 0.61 ± 0.01 | 0.24 ± 0.01 | -0.19 ± 0.05 | 2.48 ± 0.04 |
|  | Drug Average | 2.303 | 0.877 | 0.634 | N/A | -0.010 | 2.334 |
|  | DeepTTA-DB | **1.71 ± 0.02** | 0.898 ± 0.001 | **0.725 ± 0.004** | 0.507 ± 0.007 | 0.23 ± 0.01 | **1.73 ± 0.03** |
|  | DeepTTA | 1.72 ± 0.02 | **0.8991 ± 0.0008** | 0.720 ± 0.003 | **0.514 ± 0.006** | **0.236 ± 0.006** | 1.74 ± 0.02 |
| 3 | tCNNS | 2.7 ± 0.1 | 0.865 ± 0.003 | 0.57 ± 0.02 | 0.15 ± 0.02 | -0.17 ± 0.03 | 2.69 ± 0.07 |
|  | GraphDRP | 2.7 ± 0.2 | 0.872 ± 0.002 | 0.56 ± 0.03 | 0.22 ± 0.02 | -0.17 ± 0.09 | 2.7 ± 0.1 |
|  | Drug Average | 2.546 | 0.877 | 0.602 | N/A | -0.018 | 2.505 |
|  | DeepTTA-DB | **1.679 ± 0.009** | **0.8982 ± 0.0007** | **0.7303 ± 0.0009** | **0.573 ± 0.005** | **0.312 ± 0.004** | **1.676 ± 0.008** |
|  | DeepTTA | 1.70 ± 0.01 | 0.897 ± 0.001 | 0.724 ± 0.003 | **0.573 ± 0.004** | 0.305 ± 0.008 | 1.70 ± 0.01 |

**Table 11:** Cancer blind testing with no stratification (apart from last col MSE drug strat, which is only marginally different from the non-stratfied metric due to a different number of cell lines being evaluated for each drug caused by missing truth values). For three train test splits.

| Train test split | Method | MSE | Pear | R2 | MSE<br>drug strat |
| --- | --- | --- | --- | --- | --- |
| 1 | tCNNS | 2.42 ± 0.04 | 0.819 ± 0.004 | 0.660 ± 0.006 | 2.44 ± 0.03 |
|  | GraphDRP | 2.42 ± 0.08 | 0.827 ± 0.003 | 0.66 ± 0.01 | 2.43 ± 0.08 |
|  | Drug Average | 2.191 | 0.832 | 0.692 | 2.189 |
|  | DeepTTA-DB | **1.57 ± 0.01** | **0.883 ± 0.001** | **0.779 ± 0.002** | **1.57 ± 0.01** |
|  | DeepTTA | **1.57 ± 0.02** | **0.883 ± 0.002** | **0.779 ± 0.003** | **1.57 ± 0.02** |
| 2 | tCNNS | 2.412 ± 0.009 | 0.818 ± 0.002 | 0.664 ± 0.001 | 2.45 ± 0.03 |
|  | GraphDRP | 2.47 ± 0.05 | 0.821 ± 0.002 | 0.657 ± 0.007 | 2.48 ± 0.04 |
|  | Drug Average | 2.312 | 0.824 | 0.678 | 2.334 |
|  | DeepTTA-DB | **1.72 ± 0.02** | 0.873 ± 0.001 | **0.760 ± 0.003** | **1.73 ± 0.03** |
|  | DeepTTA | 1.73 ± 0.02 | **0.874 ± 0.001** | 0.759 ± 0.002 | 1.74 ± 0.02 |
| 3 | tCNNS | 2.7 ± 0.1 | 0.801 ± 0.006 | 0.64 ± 0.01 | 2.69 ± 0.07 |
|  | GraphDRP | 2.7 ± 0.1 | 0.815 ± 0.003 | 0.64 ± 0.02 | 2.7 ± 0.1 |
|  | Drug Average | 2.523 | 0.812 | 0.658 | 2.505 |
|  | DeepTTA-DB | **1.674 ± 0.009** | **0.880 ± 0.001** | **0.773 ± 0.001** | **1.676 ± 0.008** |
|  | DeepTTA | 1.70 ± 0.01 | 0.8790 ± 0.0008 | 0.770 ± 0.002 | 1.70 ± 0.01 |

Table 12 shows the results for tCNNS and GraphDRP but by replacing the genomics cell line profiles with transcriptomics profiles, (tCNNS_Tran an GraphDRP_Tran). It also shows these results for tCNNS_Tran-DB and GraphDRP_Tran-DB, removing the drug branch from the respective models. Where these tables show that adding transcriptomics cell line profiles causes the models to outperform the drug average benchmark. Furthermore, removing the drug features does not decrease performance.

**Table 12:** Cancer blind cell line stratified metrics for tCNNS and GraphDRP but by replacing the genomics cell line profiles with transcriptomics profiles, (tCNNS_Tran an GraphDRP_Tran) and for CNNS_Tran-DB and GraphDRP_Tran-DB, removing the drug branch from the respective models.

| Train test split | Method | MSE | Pear | R2 |
| --- | --- | --- | --- | --- |
| 1 | tCNNS | $2.41 \pm 0.04$ | $0.870 \pm 0.003$ | $0.61 \pm 0.01$ |
| | GraphDRP | $2.41 \pm 0.08$ | $0.872 \pm 0.004$ | $0.60 \pm 0.01$ |
|  | Drug Average | 2.190 | 0.884 | 0.640 |
| | tCNNS_Trans | $1.83 \pm 0.05$ | $0.8865 \pm 0.0005$ | $0.698 \pm 0.008$ |
| | tCNNS_Trans-DB | $1.78 \pm 0.04$ | $0.8919 \pm 0.0009$ | $0.702 \pm 0.007$ |
| | GraphDRP_Trans | $1.77 \pm 0.02$ | $0.898 \pm 0.002$ | $0.714 \pm 0.004$ |
| | GraphDRP_Trans-DB | $1.56 \pm 0.01$ | $0.9062 \pm 0.0008$ | $0.743 \pm 0.003$ |
| 2 | tCNNS | $2.4 \pm 0.005$ | $0.864 \pm 0.002$ | $0.620 \pm 0.004$ |
| | GraphDRP | $2.45 \pm 0.05$ | $0.867 \pm 0.002$ | $0.61 \pm 0.01$ |
|  | Drug Average | 2.303 | 0.877 | 0.634 |
| | tCNNS_Trans | $2.05 \pm 0.05$ | $0.875 \pm 0.003$ | $0.67 \pm 0.01$ |
| | tCNNS_Trans-DB | $1.94 \pm 0.04$ | $0.884 \pm 0.001$ | $0.685 \pm 0.007$ |
| | GraphDRP_Trans | $1.87 \pm 0.06$ | $0.898 \pm 0.001$ | $0.70 \pm 0.01$ |
| | GraphDRP_Trans-DB | $1.71 \pm 0.03$ | $0.9001 \pm 0.0007$ | $0.723 \pm 0.004$ |
| 3 | tCNNS | $2.7 \pm 0.1$ | $0.865 \pm 0.003$ | $0.57 \pm 0.02$ |
| | GraphDRP | $2.7 \pm 0.2$ | $0.872 \pm 0.002$ | $0.56 \pm 0.03$ |
|  | Drug Average | 2.546 | 0.877 | 0.602 |
| | tCNNS_Trans | $2.07 \pm 0.06$ | $0.873 \pm 0.002$ | $0.67 \pm 0.01$ |
| | tCNNS_Trans-DB | $2.00 \pm 0.02$ | $0.883 \pm 0.001$ | $0.676 \pm 0.005$ |
| | GraphDRP_Trans | $1.93 \pm 0.07$ | $0.892 \pm 0.003$ | $0.69 \pm 0.01$ |
| | GraphDRP_Trans-DB | $1.75 \pm 0.01$ | $0.8968 \pm 0.0006$ | $0.718 \pm 0.003$ |

Table 13 shows the results for first scaling the truth values to be between zero and one using the same method as in [13] before re-training and testing the models. The results agree with using the original method of pre-processing the repose values (log scaling). Specifically, they show that the null drug average benchmark outperforms the models that use genomics profiles and the models that use transcriptomics features outperform the benchmark. Furthermore, removing the drug features does not decrease the performance of DeepTTA.

**Table 13:** Cell line stratified cancer blind testing for models trained and tested with response values scaled between zero and one. The drug average benchmark outperforms the models that use genomics profiles and the models that use transcriptomics features outperform the benchmark. Furthermore, removing the drug features does not decrease the performance of DeepTTA.

|  | MAE cl strat | Pear cl strat | R2 cl strat | Pear drug strat | R2 drug strat |
| --- | --- | --- | --- | --- | --- |
| tCNNS | $0.0291 \pm 0.0001$ | $0.861 \pm 0.005$ | $0.578 \pm 0.003$ | $0.152 \pm 0.004$ | $-0.24 \pm 0.01$ |
| GraphDRP | $0.0293 \pm 0.0001$ | $0.86 \pm 0.01$ | $0.59 \pm 0.02$ | $0.161 \pm 0.008$ | $-0.25 \pm 0.03$ |
| Drug Average | 0.027 | 0.885 | 0.642 | N/A | -0.013 |
| DeepTTA-DB | **$0.0225 \pm 0.0002$** | **$0.906 \pm 0.002$** | **$0.7503 \pm 0.0009$** | **$0.538 \pm 0.002$** | **$0.274 \pm 0.004$** |
| DeepTTA | $0.0227 \pm 0.0001$ | $0.905 \pm 0.001$ | $0.746 \pm 0.005$ | $0.524 \pm 0.009$ | $0.26 \pm 0.01$ |

### I Mixed set testing

This section shows the tables for mixed set testing for additional train test splits for continuous drug response values. Note that all mixed set testing metrics calculated here were non stratified, so calculated once for all model predictions.

This section also shows the results for the marker versions of each of the literature models in Table 15. Here we removed the drug branches of the models and instead simply fed in a one-hot encoded marker representation of the drugs as we did when creating DeepTTA-DB. We also replaced the omics inputs of the models with a one-hot encoded marker representation before re-training and testing the models.

**Table 14:** Non stratified mixed set testing for train test splits 2 and 3.

| Train test split | Method | MSE | Pear | R2 |
| --- | --- | --- | --- | --- |
| 2 | tCNNS | $1.25 \pm 0.01$ | $0.9114 \pm 0.0007$ | $0.830 \pm 0.001$ |
| | GraphDRP | $1.08 \pm 0.02$ | $0.9312 \pm 0.0006$ | $0.854 \pm 0.003$ |
| | DeepTTA | $0.973 \pm 0.009$ | $0.9317 \pm 0.0008$ | $0.868 \pm 0.001$ |
|  | marker benchmark | **$0.890 \pm 0.001$** | **$0.9375 \pm 0.0001$** | **$0.8789 \pm 0.0002$** |
| 3 | tCNNS | $1.31 \pm 0.01$ | $0.9060 \pm 0.0007$ | $0.819 \pm 0.002$ |
| | GraphDRP | $1.047 \pm 0.009$ | $0.93 \pm 0.0008$ | $0.856 \pm 0.001$ |
| | DeepTTA | $1.00 \pm 0.01$ | $0.9290 \pm 0.0007$ | $0.863 \pm 0.001$ |
|  | marker benchmark | **$0.904 \pm 0.006$** | **$0.9359 \pm 0.0005$** | **$0.8757 \pm 0.0009$** |

**Table 15:**
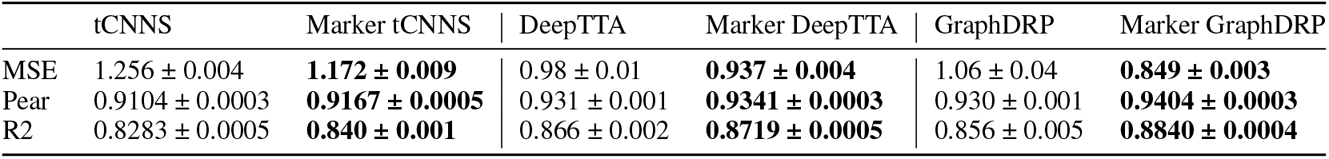
mixed set testing with maker inputs to the literature models using the original dataset. The table shows omics and drug features do not improve model performance. **Bold** gives best metric by model type.

### J Drug blind testing

This section shows the tables for non stratified drug blind testing for all three train test splits.

**Table 16:** Metrics for non-stratified drug blind testing for three train test splits. The metrics in **bold** are better than, or have bounds better than, the CL average benchmark.

| Train test split | Method | MSE | Pear | R2 |
| --- | --- | --- | --- | --- |
| 1 | GraphDRP | $11.0 \pm 3$ | $-0.0 \pm 0.2$ | $-0.5 \pm 0.4$ |
| | DeepTTA | $7.5 \pm 0.5$ | **$0.28 \pm 0.07$** | $-0.04 \pm 0.07$ |
|  | tCNNs | **$7.0 \pm 1$** | **$0.3 \pm 0.1$** | **$0.0 \pm 0.2$** |
|  | CL Average | 6.493 | 0.318 | 0.093 |
| 2 | GraphDRP | $6.0 \pm 1$ | **$0.3 \pm 0.2$** | $-0.1 \pm 0.2$ |
| | DeepTTA | $5.6 \pm 0.2$ | **$0.42 \pm 0.03$** | $-0.02 \pm 0.04$ |
| | tCNNs | $6.4 \pm 0.8$ | **$0.3 \pm 0.1$** | $-0.2 \pm 0.2$ |
|  | CL Average | 4.999 | 0.348 | 0.090 |
| 3 | GraphDRP | $11.2 \pm 0.8$ | $0.06 \pm 0.08$ | $-0.27 \pm 0.09$ |
|  | DeepTTA | **$7.7 \pm 0.6$** | **$0.47 \pm 0.01$** | **$0.13 \pm 0.07$** |
|  | tCNNs | **$6.7 \pm 0.2$** | **$0.52 \pm 0.04$** | **$0.24 \pm 0.02$** |
|  | CL Average | 8.113 | 0.288 | 0.082 |

**Table 17:**
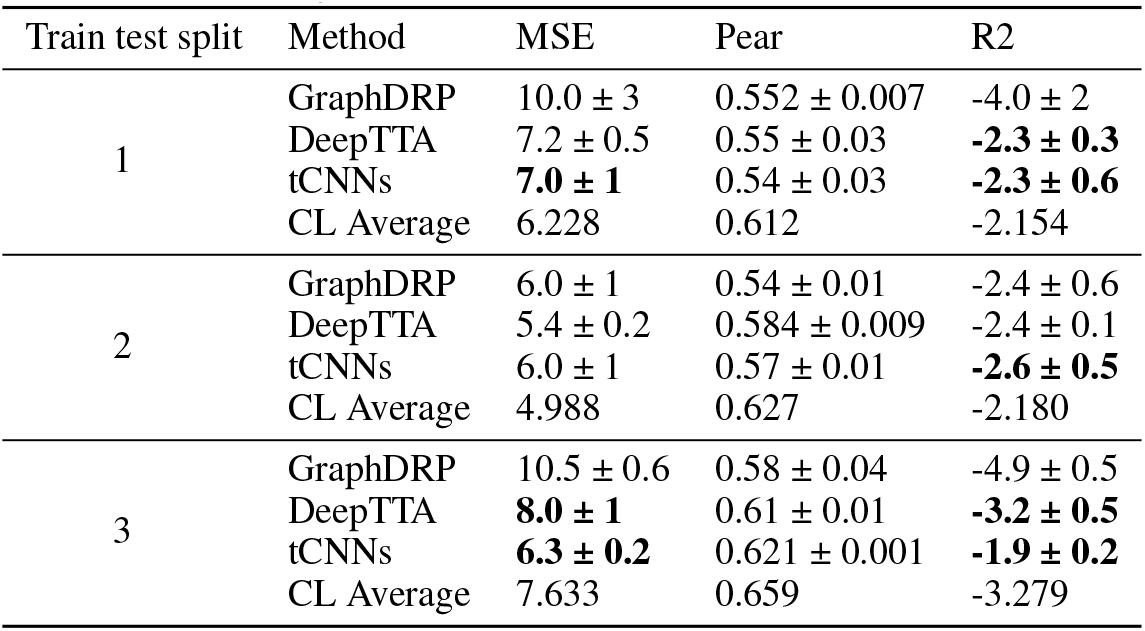
Metrics for drug stratified drug blind testing for three train test splits. The metrics in **bold** are better than, or have bounds better than, the CL average benchmark. Or better than zero for the case of Pear where the CL average benchmark is undefined.

**Table 18:** Metrics for cell line stratified drug blind testing for three train test splits. The metrics in **bold** are better than, or have bounds better than, the CL average benchmark.

| Train test split | Method | MSE | Pear | R2 |
| --- | --- | --- | --- | --- |
| 1 | GraphDRP | $11.0 \pm 3$ | $-0.2 \pm 0.2$ | **$-0.7 \pm 0.5$** |
| | DeepTTA | $7.5 \pm 0.5$ | **$0.1 \pm 0.1$** | **$-0.18 \pm 0.09$** |
|  | tCNNs | **$7.0 \pm 1$** | **$0.2 \pm 0.2$** | **$-0.1 \pm 0.2$** |
|  | CL Average | 6.503 | N/A | -0.023 |
| 2 | GraphDRP | **$6.0 \pm 1$** | **$0.1 \pm 0.2$** | **$-0.3 \pm 0.3$** |
| | DeepTTA | $5.6 \pm 0.2$ | **$0.30 \pm 0.04$** | $-0.19 \pm 0.04$ |
| | tCNNs | $6.4 \pm 0.8$ | **$0.1 \pm 0.1$** | $-0.4 \pm 0.2$ |
|  | CL Average | 5.001 | N/A | -0.057 |
| 3 | GraphDRP | $11.2 \pm 0.8$ | $-0.1 \pm 0.1$ | $-0.4 \pm 0.1$ |
| | DeepTTA | $7.7 \pm 0.6$ | **$0.407 \pm 0.008$** | **$0.04 \pm 0.08$** |
| | tCNNs | $6.7 \pm 0.2$ | **$0.47 \pm 0.06$** | **$0.17 \pm 0.03$** |
|  | CL Average | 8.125 | N/A | -0.010 |

### K Binary cancer blind testing

This section shows the tables for cancer blind testing for binary response values.

**Table 19:** Drug and cell line (CL) stratified cancer blind testing for all three train test splits, for binary response values.

| Train test Split | Method | Drug Strat<br>AUC | Drug Strat<br>AUPR | CL Strat<br>AUC | CL Strat<br>AUPR |
| --- | --- | --- | --- | --- | --- |
| 1 | Drug Average | 0.500 | 0.320 | 0.916 | 0.801 |
| | tCNNS | 0.598 ± 0.008 | 0.40 ± 0.01 | 0.905 ± 0.001 | 0.771 ± 0.005 |
| | GraphDRP | 0.6173 ± 0.0009 | 0.421 ± 0.007 | 0.911 ± 0.001 | 0.787 ± 0.002 |
| | DeepTTA-DB | 0.739 ± 0.006 | 0.53 ± 0.01 | 0.924 ± 0.001 | 0.812 ± 0.002 |
| | DeepTTA | 0.747 ± 0.008 | 0.53 ± 0.02 | 0.926 ± 0.002 | 0.814 ± 0.003 |
| | BinaryET-DB | 0.76 ± 0.01 | 0.55 ± 0.02 | 0.928 ± 0.001 | 0.817 ± 0.001 |
|  | BinaryET | **0.771 ± 0.003** | **0.569 ± 0.004** | **0.9305 ± 0.0006** | **0.822 ± 0.002** |
| 2 | Drug Average | 0.500 | 0.352 | 0.915 | 0.819 |
| | tCNNS | 0.63 ± 0.01 | 0.456 ± 0.002 | 0.9129 ± 0.0009 | 0.813 ± 0.002 |
| | GraphDRP | 0.631 ± 0.008 | 0.46 ± 0.02 | 0.912 ± 0.002 | 0.814 ± 0.004 |
| | DeepTTA-DB | 0.777 ± 0.009 | 0.60 ± 0.02 | 0.931 ± 0.003 | 0.840 ± 0.005 |
| | DeepTTA | 0.785 ± 0.002 | **0.620 ± 0.008** | 0.934 ± 0.002 | 0.845 ± 0.003 |
| | BinaryET-DB | 0.781 ± 0.009 | 0.60 ± 0.01 | 0.931 ± 0.003 | 0.839 ± 0.006 |
| | BinaryET | **0.79 ± 0.01** | 0.61 ± 0.02 | **0.934 ± 0.003** | **0.846 ± 0.006** |
| 3 | Drug Average | 0.500 | 0.339 | 0.917 | 0.817 |
| | tCNNS | 0.588 ± 0.008 | 0.427 ± 0.005 | 0.9055 ± 0.0008 | 0.785 ± 0.004 |
| | GraphDRP | 0.599 ± 0.008 | 0.43 ± 0.01 | 0.9109 ± 0.0009 | 0.803 ± 0.001 |
| | DeepTTA-DB | 0.785 ± 0.007 | 0.6 ± 0.008 | 0.9308 ± 0.0007 | 0.8390 ± 0.0005 |
| | DeepTTA | 0.789 ± 0.005 | 0.601 ± 0.003 | 0.932 ± 0.001 | 0.842 ± 0.002 |
| | BinaryET-DB | 0.795 ± 0.003 | 0.608 ± 0.002 | 0.933 ± 0.001 | 0.841 ± 0.004 |
|  | BinaryET | **0.798 ± 0.003** | **0.613 ± 0.005** | **0.9348 ± 0.0006** | **0.844 ± 0.001** |

**Table 20:** Non stratified cancer blind testing.

| Train test split | Method | AUC | AUPR |
| --- | --- | --- | --- |
| 1 | Drug Average | 0.884 | 0.746 |
| | tCNNS | 0.878 ± 0.001 | 0.735 ± 0.004 |
| | GraphDRP | 0.8830 ± 0.0009 | 0.7468 ± 0.0009 |
| | DeepTTA | 0.9101 ± 0.0008 | 0.799 ± 0.002 |
|  | BinaryET | **0.9158 ± 0.0006** | **0.8084 ± 0.0002** |
| 2 | Drug Average | 0.877 | 0.761 |
| | tCNNS | 0.879 ± 0.003 | 0.766 ± 0.001 |
| | GraphDRP | 0.881 ± 0.002 | 0.774 ± 0.003 |
|  | DeepTTA | **0.920 ± 0.002** | **0.841 ± 0.004** |
| | BinaryET | 0.919 ± 0.003 | 0.840 ± 0.005 |
| 3 | Drug Average | 0.886 | 0.765 |
| | tCNNS | 0.879 ± 0.001 | 0.757 ± 0.003 |
| | GraphDRP | 0.8829 ± 0.0006 | 0.765 ± 0.002 |
| | DeepTTA | 0.9212 ± 0.0008 | 0.837 ± 0.001 |
|  | BinaryET | **0.9239 ± 0.0005** | **0.840 ± 0.001** |

**Table 21:** Ablation study for DeepTTA for multiple split types and testing type, the table shows that removing the drug branch decreases performance. For given split and testing time the **bold** metric gives the best performance between DeepTTA and DeepTTA-DB.

| Strat type | Train test split | DeepTTA AUC | DeepTTA-DB AUC | DeepTTA AUPR | DeepTTA-DB AUC |
| --- | --- | --- | --- | --- | --- |
| Drug strat | 1 | **0.747 ± 0.008** | 0.739 ± 0.006 | **0.53 ± 0.02** | **0.53 ± 0.01** |
| | 2 | **0.785 ± 0.002** | 0.777 ± 0.009 | **0.620 ± 0.008** | 0.60 ± 0.02 |
| | 3 | **0.789 ± 0.005** | 0.785 ± 0.007 | **0.601 ± 0.003** | 0.6 ± 0.008 |
| CL strat | 1 | **0.926 ± 0.002** | 0.924 ± 0.001 | **0.814 ± 0.003** | 0.812 ± 0.002 |
| | 2 | **0.934 ± 0.002** | 0.931 ± 0.003 | **0.845 ± 0.003** | 0.840 ± 0.005 |
| | 3 | **0.932 ± 0.001** | 0.9308 ± 0.0007 | **0.842 ± 0.002** | 0.8390 ± 0.0005 |
| No strat | 1 | **0.9101 ± 0.0008** | 0.909 ± 0.002 | **0.799 ± 0.002** | **0.799 ± 0.003** |
| | 2 | **0.920 ± 0.002** | 0.917 ± 0.003 | **0.841 ± 0.004** | 0.836 ± 0.005 |
| | 3 | **0.9212 ± 0.0008** | 0.9196 ± 0.0009 | **0.837 ± 0.001** | 0.834 ± 0.001 |

### L Binary mixed set testing

This section shows the tables for mixed set testing for binary response values.

**Table 22:** Mixed set testing for train test split 2 and 3 with binary response values.

| Methods | AUC | AUPR |
| --- | --- | --- |
| Marker | 0.9286 ± 0.0003 | 0.8691 ± 0.0005 |
| tCNNS | 0.9263 ± 0.0009 | 0.864 ± 0.002 |
| GraphDRP | 0.9354 ± 0.0004 | 0.880 ± 0.002 |
| DeepTTA | 0.9406 ± 0.0009 | 0.891 ± 0.003 |
| BinaryET-DB | 0.9451 ± 0.0005 | 0.8975 ± 0.0009 |
| BinaryET | **0.946 ± 0.001** | **0.9 ± 0.002** |
| Marker | 0.9284 ± 0.0001 | 0.8648 ± 0.0003 |
| tCNNS | 0.9244 ± 0.0006 | 0.8576 ± 0.0007 |
| GraphDRP | 0.9339 ± 0.0005 | 0.875 ± 0.001 |
| DeepTTA | 0.9435 ± 0.0002 | 0.8923 ± 0.0002 |
| BinaryET-DB | 0.9455 ± 0.0007 | 0.896 ± 0.001 |
| BinaryET | **0.9466 ± 0.0007** | **0.898 ± 0.001** |

